## Supplementary Figures for "Dominant toxin hypothesis: unravelling the venom phenotype across micro and macroevolution"

### <u>Supplementary tables</u>

**Supplementary Table 1.** Sea anemone comparative transcriptomics and completeness.

**Supplementary Table 2.** Toxin expression matrix among sea anemones. Act = Actinioidea, Met = Metridioidea, Edw = Edwardsioidea

**Supplementary Table 3.** Phylogenetic covariance analysis of toxin expression among sea anemones. NS = Not significant.

**Supplementary Table 4.** Phylogenetic signal and model of evolution on toxin expression in sea anemones. Brownian motion (BM), Ornstein–Uhlenbeck (OU), or early burst (EB) models

**Supplementary Table 5.** Phylogenetic covariance analysis and fitting tissue type as a fixed effect, showing that the confidence intervals overlap zero.

**Supplementary Table 6.** Toxin copy number in sea anemone genomes identified using BLASTp with KTx3 and NaTx highlighted in cyan as those identified manually.

**Supplementary Table 7.** Macrosynteny of the number of single-copy orthologs (SCO) shared among sea anemones genomes.

**Supplementary Table 8.** NEP3 and NEP6 genome loci in sea anemones genomes

**Supplementary Table 9.** Percent identity for NaTx and KTx3 genes within sea anemones genomes.

**Supplementary Table 10.** Population transcriptomics for *Nematostella vectensis* representing sequencing depth and completeness.

**Supplementary Table 11.** Gene expression of *Nv1* and other toxins in *Nematostella vectensis* from different populations. A) A matrix of known toxin genes mapped to reference *Nematostella vectensis* from different populations B) nCounter normalized data of *Nv1* expression and housekeeping genes from different populations at different temperatures.

**Supplementary Table 12.** Raw quantification cycle values and estimated *Nv1* copy number for *N. vectensis* individuals from five populations along the Atlantic Coast of North America.

**Supplementary Table 13.** Results of two-way ANOVA testing the effect of population and plate on estimates of diploid *Nv1* copy number

**Supplementary Table 14.** Tukey HSD results for *Nv1* diploid copy number across populations. Absolute difference in mean copy number is show above the diagonal and p-value below the diagonal.

**Supplementary Table 15.** Semi-quantitative proteomic results from two Florida and North Carolina populations of *Nematostella vectensis.*

**Supplementary Table 16.** Perseus statistical analysis performed using semi-quantitative proteomic results from Florida and North Carolina populations of *Nematostella vectensis.*

**Supplementary Table 17.** Raw data used for linear regression analysis comparing the semi-quantitative proteomic results from two Florida and North Carolina populations of *Nematostella vectensis.*

**Supplementary Table 18.** Read information for reads spanning *Nv1* haplotypes.

**Supplementary Table 19.** Transposable element annotations generated by EDTA spanning the *Nv1* locus and 20kb up- and downstream for the Nova Scotia, Florida, and Maryland assemblies.

### Supplementary figures

23/6/22 1:58:00 am

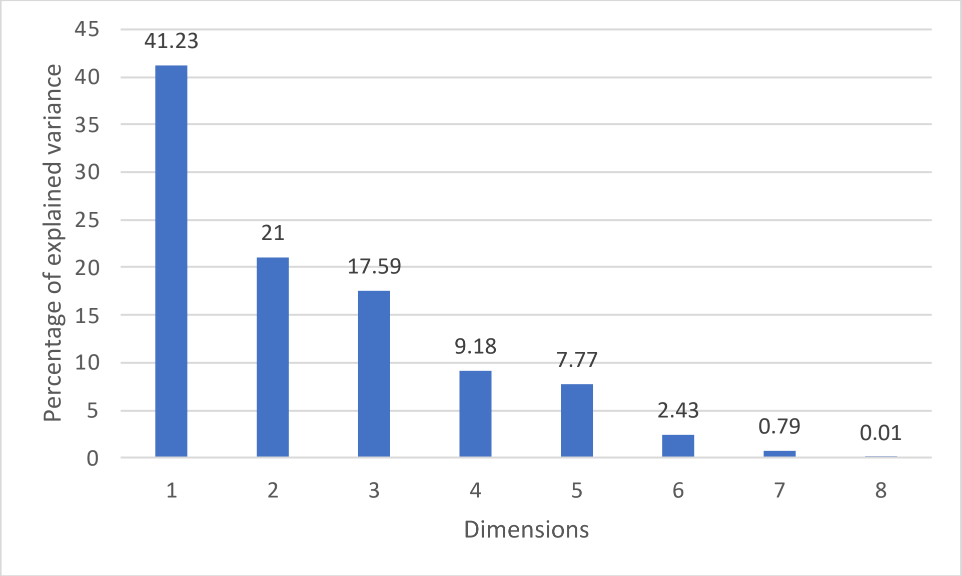


**Supplementary Figure 1.** Scree plot of the phylogenetic covariance analysis of toxin expression

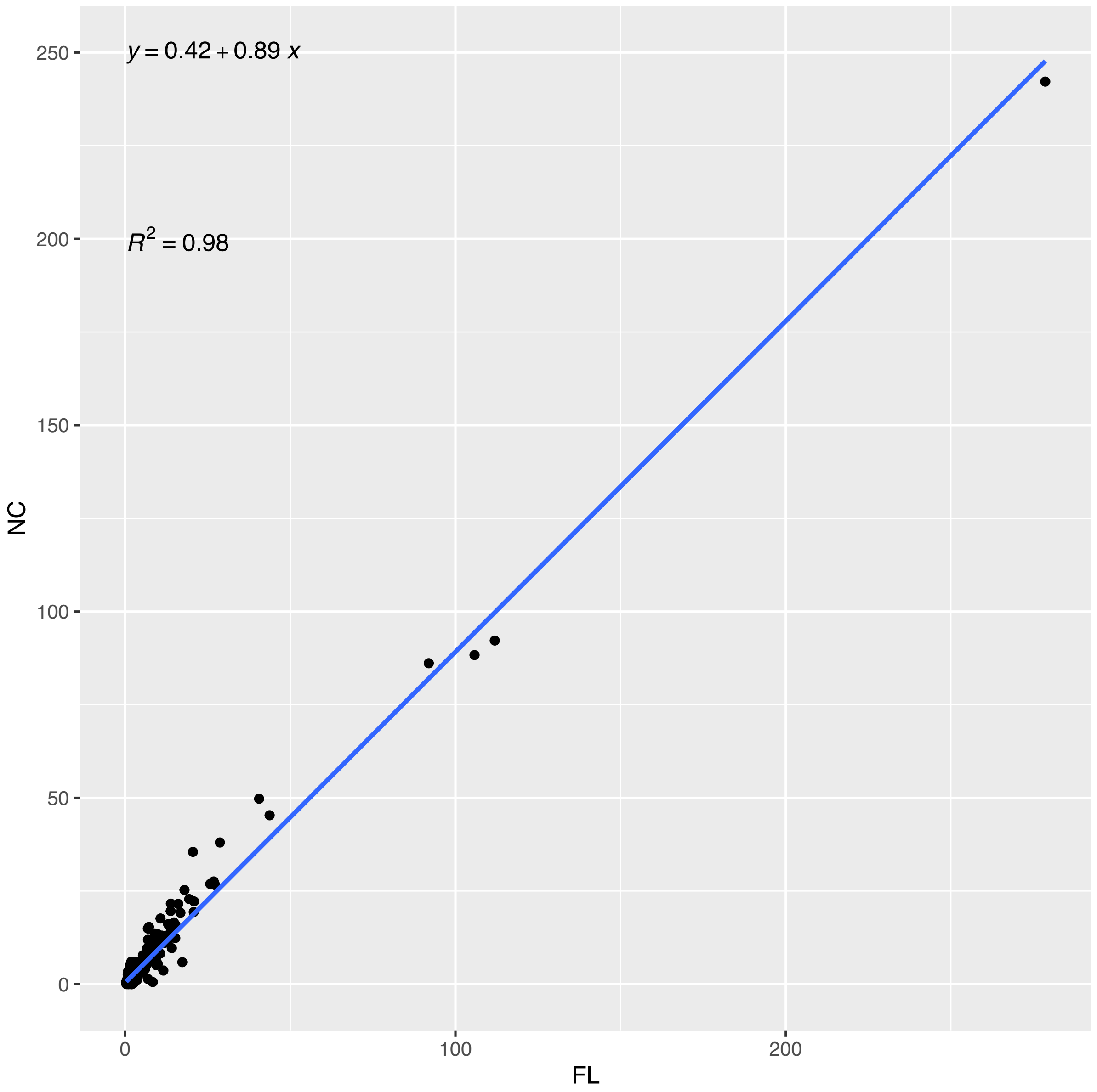


**Supplementary Figure 2.** Linear regression comparing proteomes average label-free quantification (LFQ) per million from Florida and North Carolina, on the x- and y-axis, respectively.

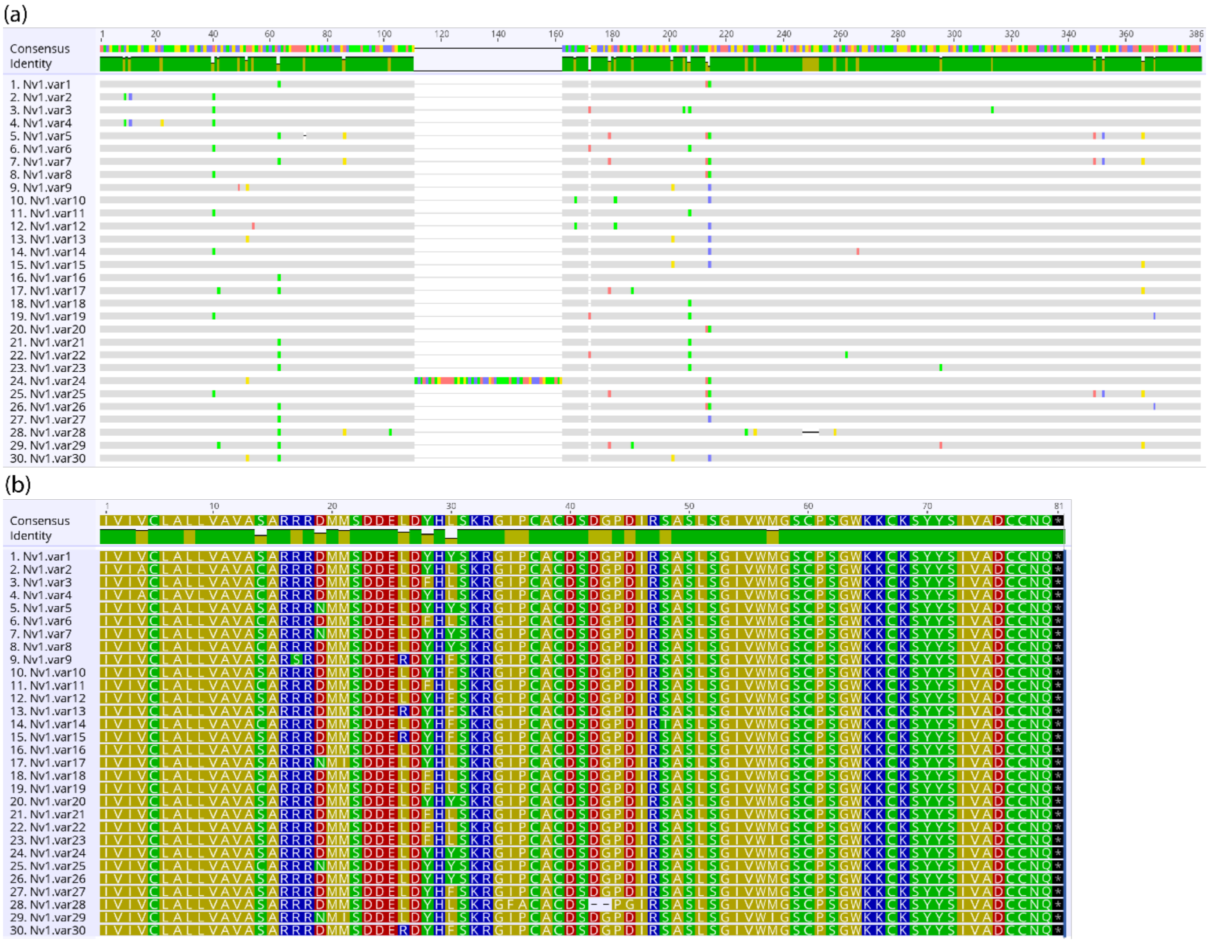


**Supplementary Figure 3.** Multiple sequence alignment of nucleotides (a) and amino acids (b) from all Nv1 paralogs identified in the amplicon analysis.

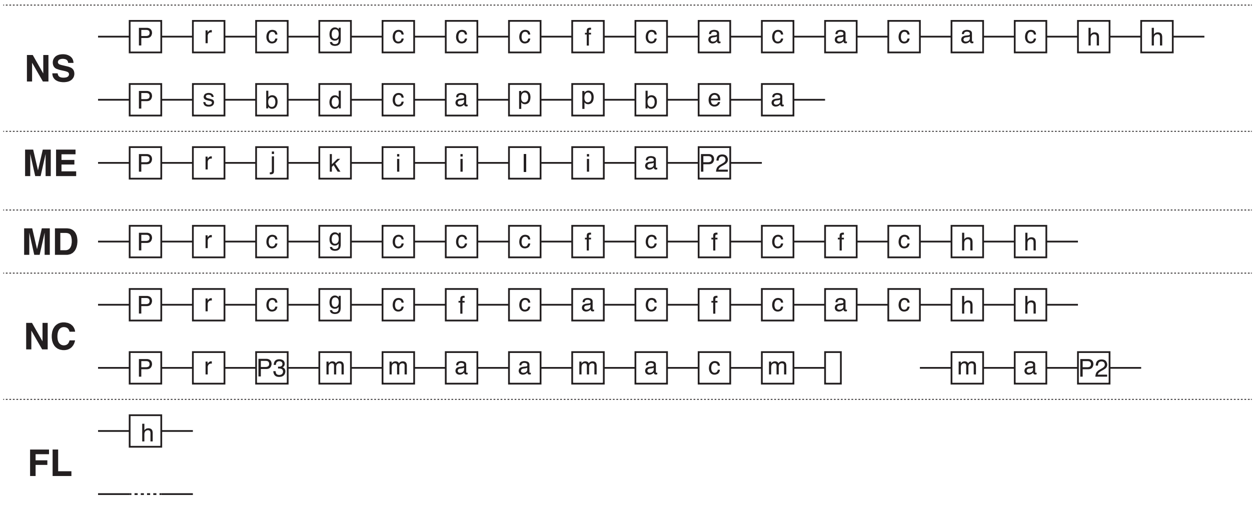


**Supplementary Figure 4.** Genomic arrangement of *Nv1* amino acid variants at the *Nv1* locus across individuals. Each letter code represents a distinct variant. Shared pseudogenes are labelled (P, P2 and P3).

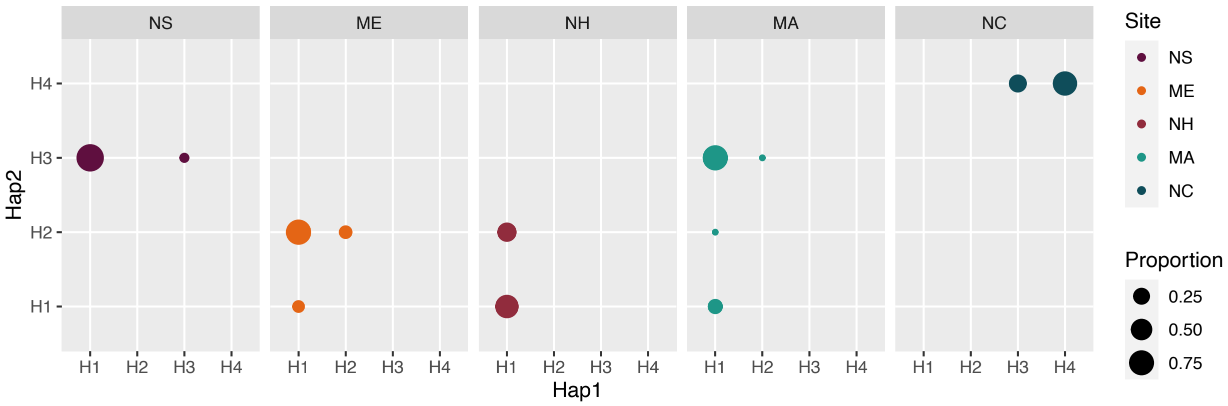


**Supplementary Figure 5.** Relative *Nv1* locus genotype frequencies between sampling sites.

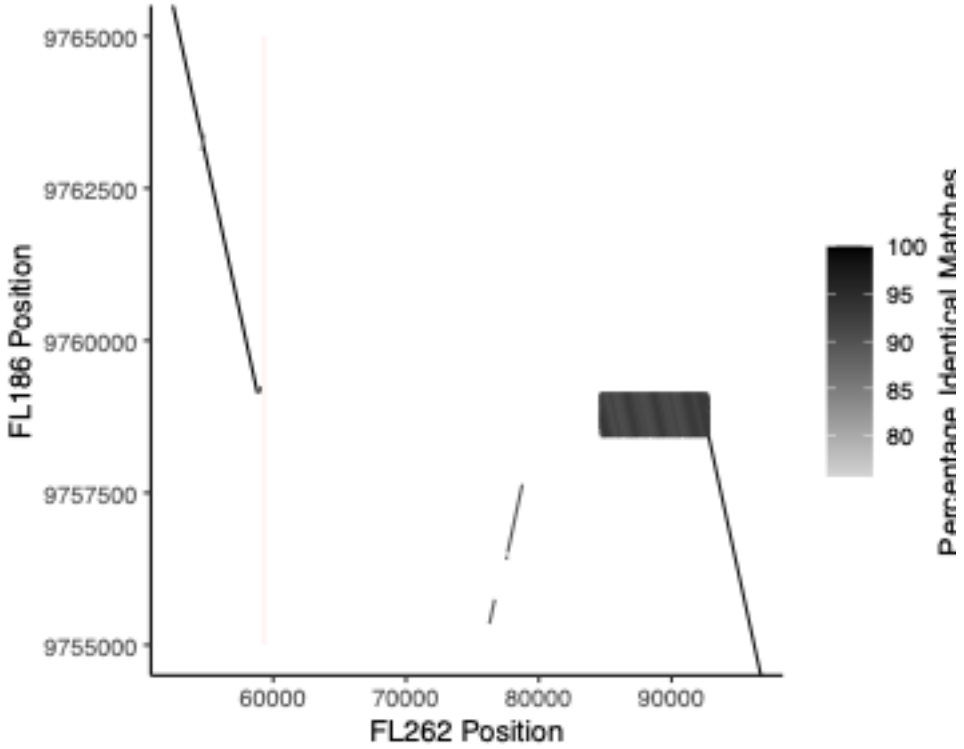


**Supplementary Figure 6.** Dotplot of BLASTn results between the two Florida haplotypes. The red bars indicate the location of the *Nv1* exons in haplotype FL262.

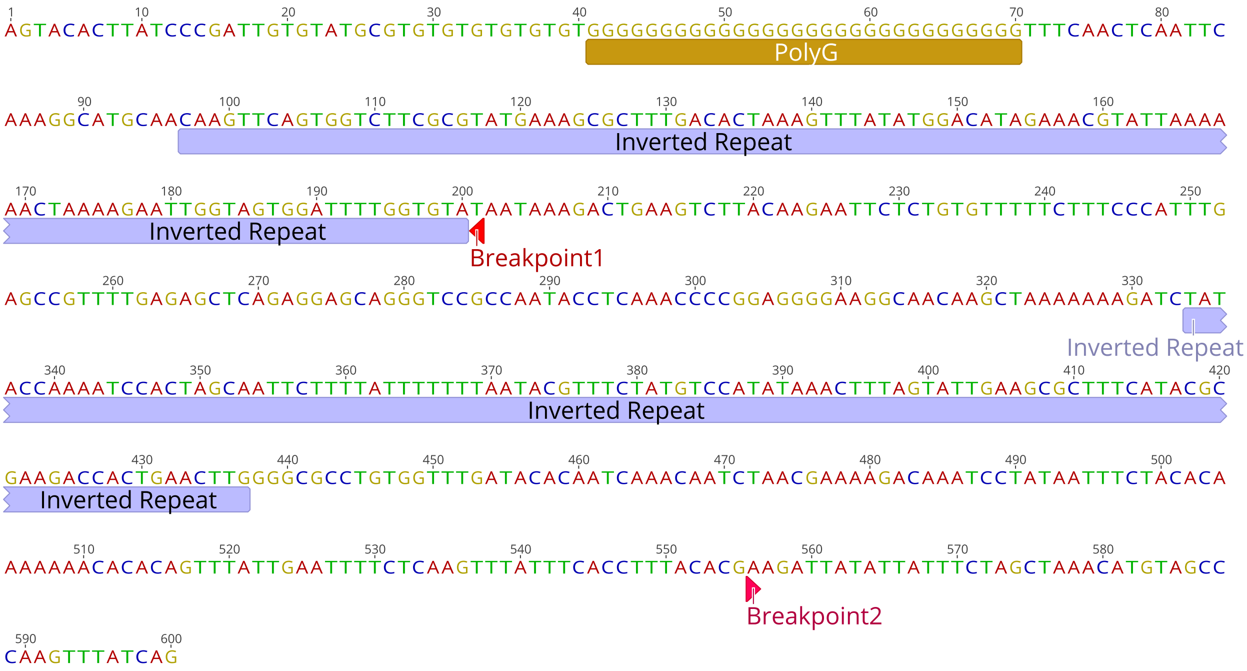


**Supplementary Figure 7.** Annotation of genomic locus (NS.tig889:951254-951854) associated with deletion breakpoints. Sequence features associated with non-b DNA structures are annotated in gold and blue. Breakpoints 1 and 2 refer to the breakpoints associated with the Nv1 locus deletion in the zero-copy haplotype, and the pseudogenes associated with NC.H4 and ME.H2, respectively.

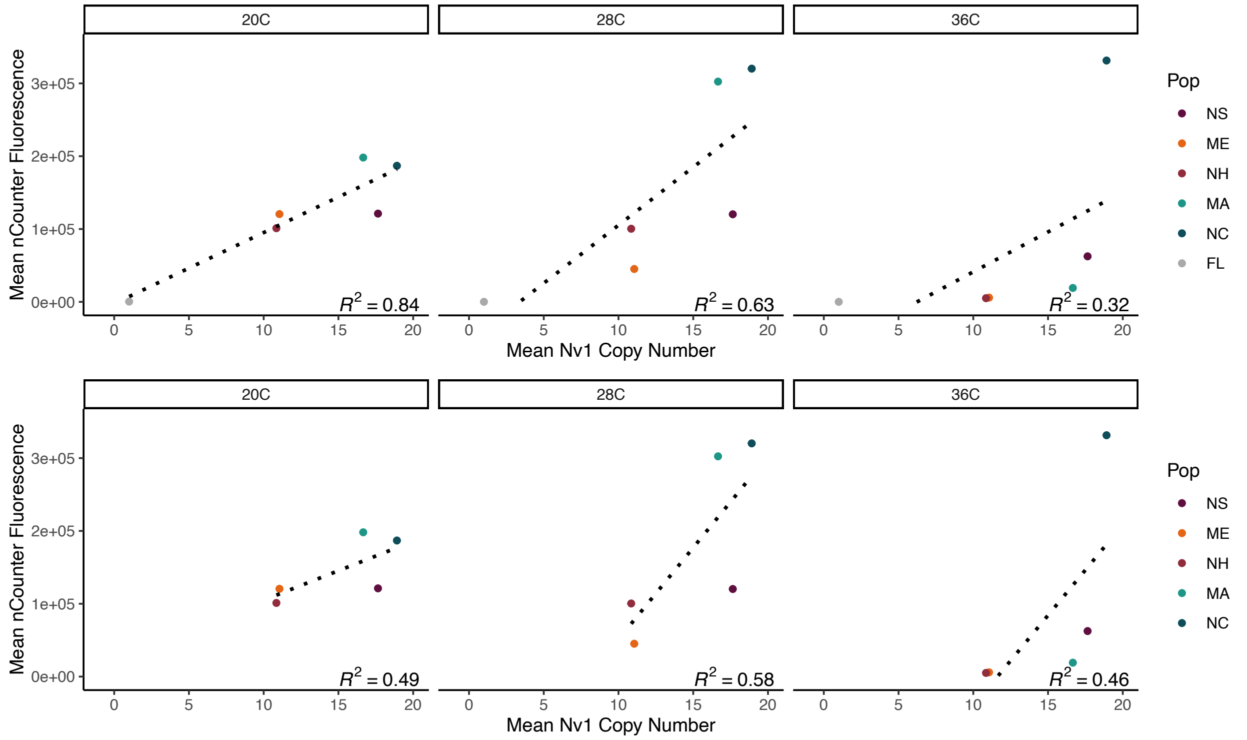


Supplementary Figure 8. Nv1 copy number vs transcript abundance at three temperatures (20°C, 28°C, and 36°C) across *N. vectensis* populations with (above) and without (below) the Florida population.

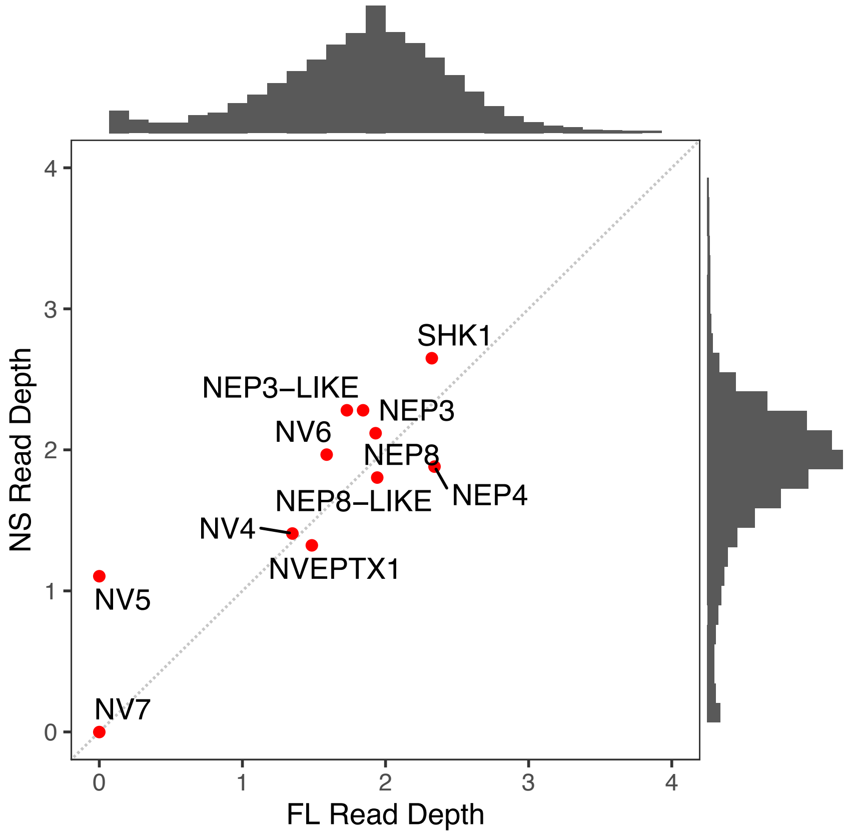

Supplementary Figure 9. Modal read depth for Nova Scotia and Florida individuals at 11 other *N. vectensis* toxin gene loci. Read depths are normalized to diploid copy number. Histograms show a representative read depth distribution sampled across 1 million bases of the *N. vectensis* genome.

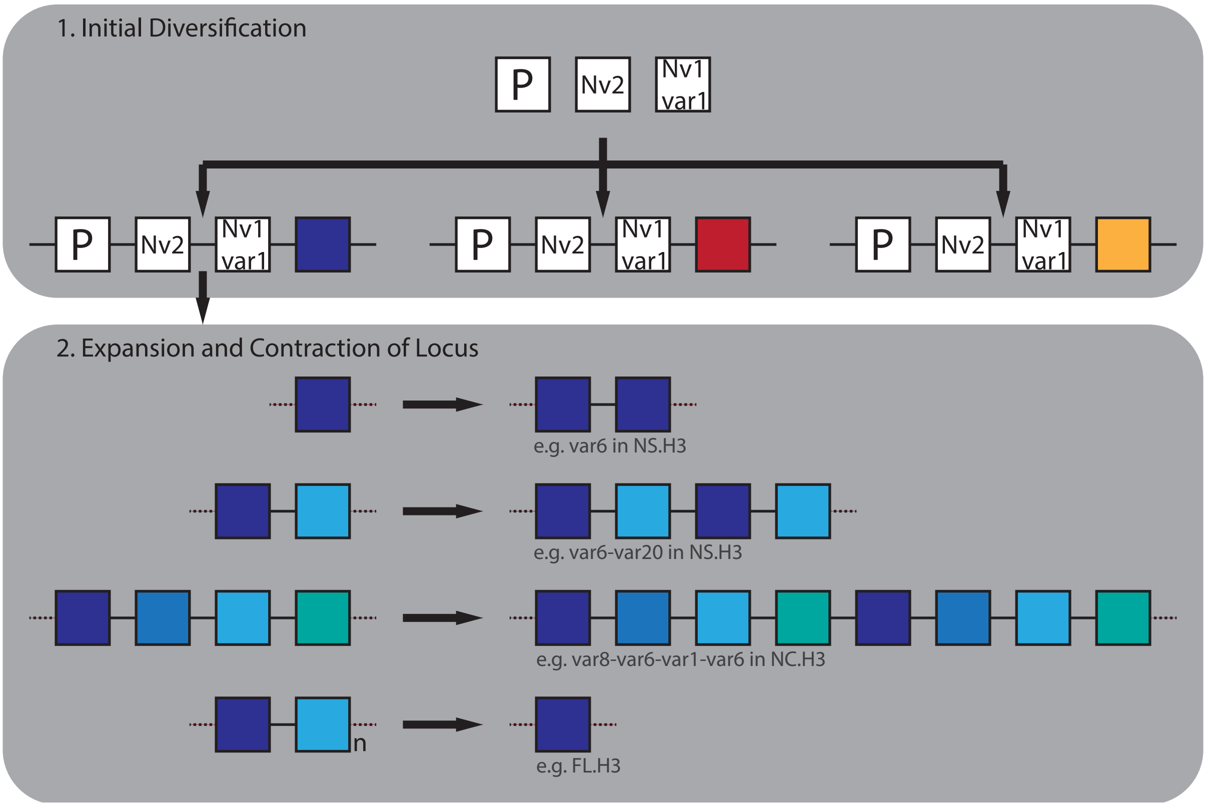


Supplementary Figure 10. Schematic representation of Nv1 locus evolution. 1) Ancestral locus (containing the pseudogene, *Nv2*, and *Nv1.var1*) undergoes initial duplication and diversification events. 2) The resulting loci subsequently expand through replication slippage generating duplicated blocks of single or multiple genes. These loci are also subject to contraction events that appear to be mediated by deletions associated with non-b DNA structures.
